# Deep Learning disconnectomes to accelerate and improve long-term predictions for post-stroke symptoms

**DOI:** 10.1101/2023.09.12.557396

**Authors:** Anna Matsulevits, Pierrick Coupe, Huy-Dung Nguyen, Lia Talozzi, Chris Foulon, Parashkev Nachev, Maurizio Corbetta, Thomas Tourdias, Michel Thiebaut de Schotten

**Affiliations:** Groupe d’Imagerie Neurofonctionnelle, Institut des Maladies Neurodégénératives-UMR 5293, CNRS, CEA, University of Bordeaux, Bordeaux, France; Brain Connectivity and Behaviour Laboratory, Sorbonne Universities, Paris, France; University Bordeaux, CNRS, Bordeaux INP, LaBRI, UMR 5800, 33400, Talence, France; Department of Neurology and Neurological Sciences, Stanford University School of Medicine, Stanford, CA, 94305, USA; Institute of Neurology, UCL, London, UK; Clinica Neurologica, Department of Neuroscience, University of Padova, Padova, Italy; Padova Neuroscience Center (PNC), University of Padova, Padova, Italy; Venetian Institute of Molecular Medicine, VIMM, Padova, Italy; Inserm-U1215, Neurocentre Magendie; CHU de Bordeaux, Neuroimagerie diagnostique et thérapeutique, Bordeaux France

**Author notes:** Authors contributed equally (Thomas Tourdias, Michel Thiebaut de Schotten).

**Keywords:** Deep Learning, Deep-Disconnectome, Stroke, Precision Medicine, Neuroimaging, White Matter, Artificial Intelligence

## Abstract

Deep learning as a truly transformative force is revolutionizing a wide range of fields, making a significant difference in medical imaging, where recent advancements have yielded some truly remarkable outcomes. In a connected brain, maps of white matter damage — otherwise known as disconnectomes — are essential for capturing the effects of focal lesions. However, the current tools for obtaining such information are prohibitively slow and not admitted for clinical usage. Here, we have explored the potential of deep-learning models to accurately generate disconnectomes in a population of stroke survivors. We trained a 3D U-Net algorithm to produce *deep-disconnectomes* from binary lesion masks. This artificial neural network was able to capture most information obtained in conventional disconnectomes, i.e., statistical maps filtering normative white-matter networks, but output a deep-disconnectome 170 times faster – compared to disconnectome computation with the state-of-the-art BCBToolkit software. Moreover, the deep-disconnectomes were challenged to predict cognitive and behavioral outcomes one-year post-stroke. In an additional cohort of N=139 stroke survivors, N=86 neuropsychological scores were predicted from deep-disconnectomes achieving, on average, 85.2% of accuracy and R^2^= 0.208. The deep-disconnectomes predictivity power outperformed the conventional disconnectome predictions for clinical scores.

In summary, we have achieved a significant milestone for clinical neuroimaging by accelerating and ameliorating the creation of disconnectome maps using deep learning. By integrating deep learning into the management of stroke, one of the most prevailing catalysts for acquired disabilities, we deepen our understanding of its impact on the brain. This novel approach may offer potential avenues for acute intervention, ultimately enhancing patients’ overall quality of life.

## 1. Introduction

Does deep learning hold promises for an improvement of society through faster and more accurate access to results, or will it degrade our scientific progress? Healthcare is exemplary for where remarkable developments regarding the inclusion of computer science technologies have been achieved, yielding fruitful outcomes and attracting globally increasingly more investments for projects involving artificial intelligence (AI) (Mou, 2019). In the past decades, deep learning models have been a particularly great addition to the field of medical imaging (Alexander et al., 2020; Panayides et al., 2020). The very first application of a deep learning model in healthcare can be traced back to 1995, when a convolutional neural network was successfully used to detect lung nodules in chest X-rays (Lo et al., 1995). Focusing on the virtual branch of artificial intelligence that represents different types of machine learning and deep learning where algorithms improve by updating their performance from experience (Hamet & Tremblay, 2017), there are over 500 FDA-approved artificial intelligence-powered algorithms and devices that are currently in use in real-life clinical settings (U.S. Food and Drug Administration, 2022). The majority (>75%) of these are devices for radiology, followed by cardiology (11%), hematology (3%), and neurology (3%). With radiology being based on imaging technologies, the field has demonstrated some highly promising developments in tasks such as image classification, semantic segmentation, instance segmentation, and image reconstruction (Cheng et al., 2021). All these challenges that are being solved with the help of artificial intelligence models in radiology are closely related to imaging in neuroscientific research and have already been impactfully applied in both fields to improve and fasten processes, such as brain tumor and lesion segmentation (Ballestar & Vilaplana, 2021; Komnitsas et al., 2017), and enhancement of MR images (Rudie et al., 2022).

In the context of neuroimaging, the last two decades have pushed forward the role of connections in the functional organization of the brain and the neurological symptoms in patients (Thiebaut de Schotten & Forkel, 2022; Fox, 2018). Whole brain connections – tractograms – can be reconstructed with diffusion MRI (Magnetic Resonance Imaging) that estimates the direction of microscopic water molecules diffusion that preferentially follows the direction of the fibers (Basser, Mattiello & LeBihan, 1994; Conturo et al., 1999; Jones et al., 1999, Mori & Barker, 1999). The acquisition of diffusion MRI is now widely feasible on modern MRI scanners and clinical settings. However, many parameters need to be set to obtain diffusion scans suitable for reconstructing white matter (WM) tracts. Moreover, the virtual dissection of WM tracts is often time-consuming and requires trained personnel. For these reasons, WM investigations are rarely integrated into clinical settings, which, especially for stroke patients, requires prompt solutions to the “time is brain” emergency answer.

A first answer in providing faster and more applicable white matter reconstructions was to exploit tractograms of normative populations to estimate the probability of disconnections given a certain lesion in the brain. Accordingly, brain disconnection maps — disconnectomes — are derived from patients’ binary lesions, which could be easily segmented from clinical MRI scans conventionally acquired in hospitals (Foulon, 2018; Kucuyeski, 2013; Griffith, 2021). Such brain disconnection patterns computed a few days after stroke showed accurate predictions (average accuracy > 80%) of the long-term impairment on different neuropsychological tests (Talozzi et al., 2023). Obtaining such detailed predictions could be highly beneficial, especially in acute clinical settings, to personalize the care pathway. However, while solving some problems related to data acquisition for tractography, the current state-of-the-art in disconnectome analyses present issues regarding calculation time and, in particular, memory space. Hence, obtaining the disconnectome from any toolkit is still a resource-consuming process (Foulon et al., 2018; Kuceyeski et al., 2013; Griffis et al., 2021), which makes it less applicable in clinical settings where time is a critical factor. Yet, it has never been tested whether deep learning could potentially make the complex and accurate 3D reconstruction of disconnected fibers a fast and automatic process while preserving clinical prediction accuracy.

To achieve this goal, we created and evaluated a 3D U-Net deep learning model to predict *deep-disconnectomes* for lesioned brains. Detailed descriptions of the used datasets are summarized in the supplementary material in Table 1. We created a 3D U-Net model (Ronnenberger, Fischer & Brox, 2015) that consists of 4 encoding- and 4 decoding layers and trained the model on 1333 synthetically created lesion-disconnectome dataset pairs (dataset 1). After training and validating the model, we tested it on an out-of-sample dataset of 1333 natural stroke lesions (dataset 2). Additionally, we tested the predictive power of deep-disconnectomes to predict clinical scores following a recently published approach on datasets 3 and 4 (Talozzi et al., 2023). The accuracy of the neuropsychological predictions derived from the deep-disconnectomes was better than conventional disconnectomes (*t*(85) = -1.663, *p* = .009), further highlighting the advances that deep learning approaches can bring to the clinic.

## 2. Results

### 2.1 Accuracy in *deep-disconnectome* reconstruction

Training the model for 260 epochs took 33.527 hours on the NVIDIA Quadro P4000 GPU. The resulting training loss and validation loss were 0.01652 and 0.01767, respectively (Figure 3b). After training the model, the average time required to generate a deep-disconnectome from a binary lesion mask was 0.642 seconds, which is significantly faster compared to the conventional disconnectome generation time of 480 seconds/8 minutes (on a standard computer setup with an Intel Core i7 processor).

Figure 1a shows an out-of-sample example of the conventional and deep-learning version of the disconnectome for the same subject. The overall goodness of fit metric, R^2^, reached 0.82, meaning that the 3D U-Net model is able to explain, on average, more than 80% of the variance in the predicted image based on the input data. Visual comparison of the predicted and the ground truth (conventional) disconnectomes demonstrates the reliability of the predictions to capture the relevant features and resemble the conventional disconnectome (Figure 1a). Running a paired t-test between conventional disconnectomes and the corresponding deep-disconnectomes further reveals the disconnected structures that are reliably captured alongside the less well-captured structures. The t-map visualization (Figure 1b) demonstrates significantly higher values for the core long white matter structures and lower values, hence an underestimation of short, U-shaped association fibers located near the surface.

**Figure 1:**
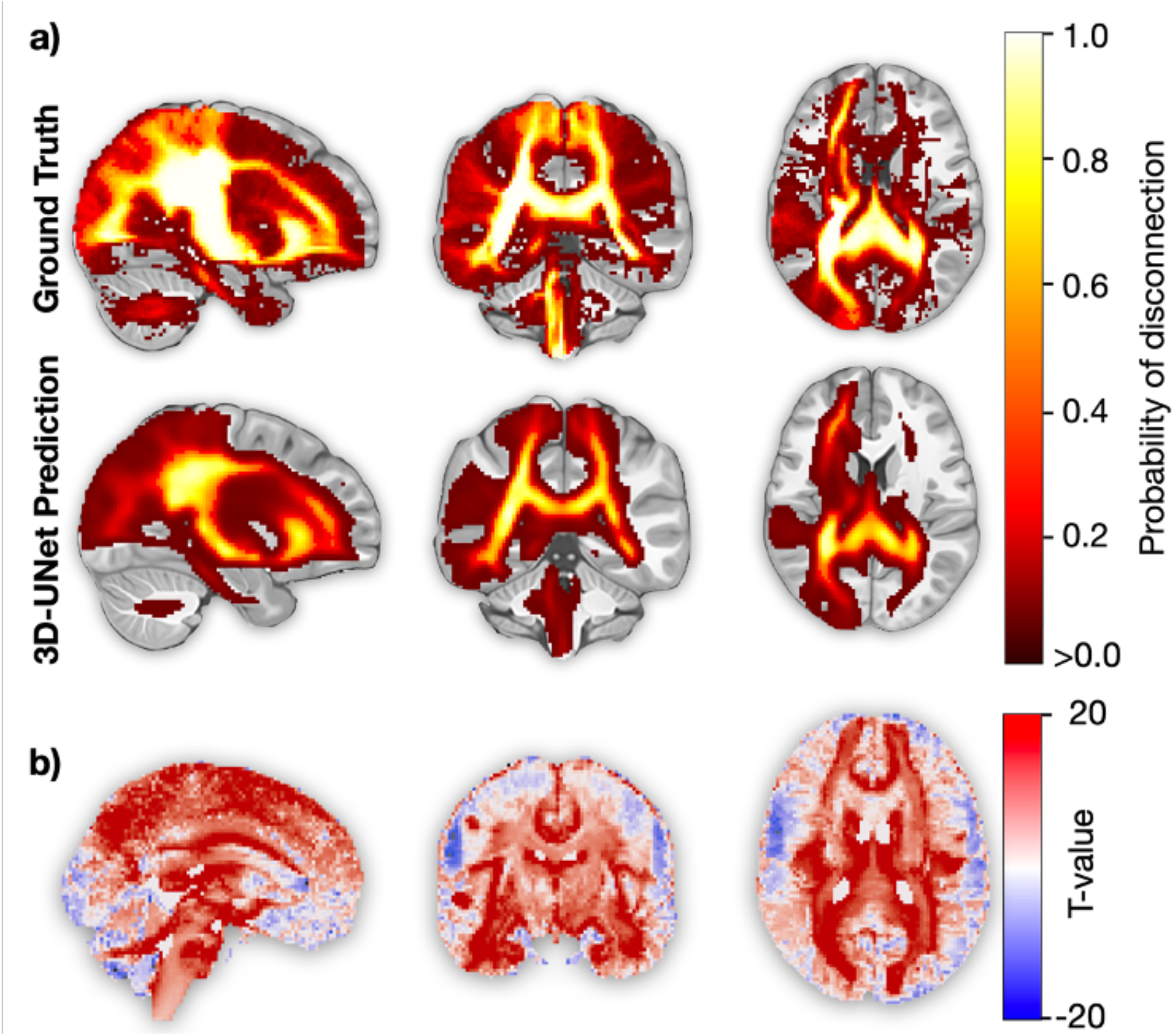
(a) Visual comparison of a representative out-of-sample patient conventional disconnectome (top) with its 3D U-Net predicted deep-disconnectome (bottom). (b) T-map obtained from the paired t-test comparing 1333 conventional and deep-disconnectomes (dataset 2). Positive t-values (red) indicate more frequent disconnections in the deep-disconnectome compared to the ground truth, while negative t-values (blue) indicate less frequent disconnections.

### 2.2 Deep-disconnectome predictivity power for neuropsychological scores

Based on the work of Talozzi et al. (2023), we used the same N=1333 stroke lesions (dataset 2) to derive deep-disconnectomes. The same dataset was imported into the Uniform Manifold Approximation and Projection (UMAP) embedding (McInnes, Healy & Melville, 2018), obtaining a novel morphospace for deep-disconnectomes (Figure 2a).

To test the deep-disconnectome predictions of neuropsychological scores and compare these to the predictions of the conventional disconnectomes, we embedded a dataset of new patients (n = 119, dataset 3) into the novel morphospace and followed the analysis steps described in Talozzi et al. (2023). With this method, we were able to predict N=86 different behavioral and cognitive scores. An average accuracy of 85.2% and an average R^2^= 0.208 were achieved. For comparison, the average accuracy and R^2^ for the exact same predictions with a morphospace obtained using disconnectomes computed by the BCBToolkit software reached 83.7% and R^2^ = 0.191, respectively. Hence, computing the disconnectomes using the deep-learning model and obtaining a corresponding morphospace for predicting neuropsychological scores significantly increases the accuracy of the resulting scores by at least 1.5%; (*t*(85) = -1.663, *p* = .009) (Figure 2b).

To perform an out-of-sample validation, N=20 stroke survivors from dataset 3 were excluded from the prediction training phase and used as independent model validation. We outputted the deep-disconnectomes from the segmented binary stroke lesion (as reported in 4.2) and predicted their corresponding neuropsychological scores (as reported in 4.4). The out-of-sample predictions achieved an average accuracy of 80.67%, with an R^2^=0.143. In comparison, the previous attempt with conventional disconnectomes (Talozzi et al., 2023) reached an average accuracy of 80.35% with a lower average R^2^ = 0.121 (Figure 1, Supplementary Material).

**Figure 2.**
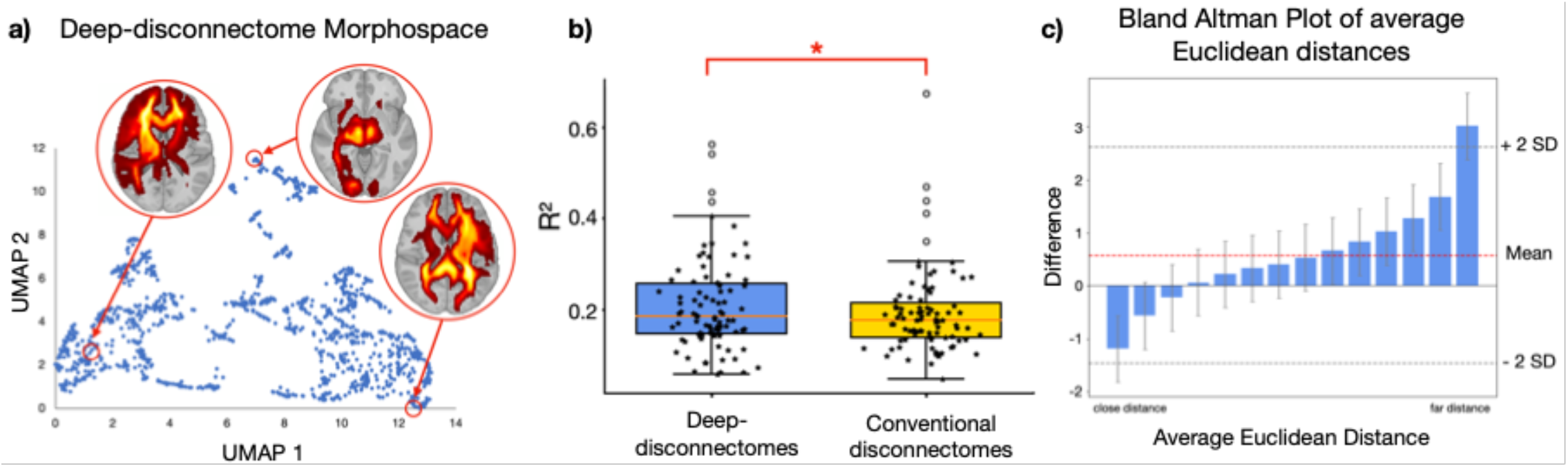
(a) UMAP morphospace embedding of N=1333 deep-disconnectomes with three exemplary deep-disconnectomes in different locations of the latent space. (b) The boxplot shows all the R^2^ for the predictions for stroke survivors (dataset 3) across N=86 neuropsychological scores for the framework using deep-disconnectomes compared to the framework using disconnectomes obtained by BCBToolkit. The boxes represent the quartiles, the whiskers indicate the distribution, and the outliers are marked as white dots. Inside the boxes, the median is visualized by the red line. The *P*-value obtained from a paired *t*-test (2-tails): \**P* < 0.01 shows a significant difference. (c) The Bland Altman plot shows the difference in the average Euclidean distances between the conventional disconnectome and the deep-disconnectome for each of the 139 disconnectomes (dataset 3 and dataset 4) to the rest of the data points in their given morphospace. Deep-disconnectomes demonstrate a better dissociation within the clusters (for close distances), as indicated by the negative differences. For far distances of the data points in the morphospace, deep-disconnectomes are located closer to each other than conventional disconnectomes, as indicated by the positive difference value. The error bars represent 95%-confidence intervals.

In order to understand why the score prediction of the deep-disconnectome outperforms the conventional disconnectome, a systematic comparison across the embedding coordinates was performed for each patient. We assessed the structure of the latent space by means of the average Euclidean distance between each embedding set of coordinates (UMAP 1 and 2) against the rest of the embedding points. This calculation reveals a higher distance within clusters of the deep-disconnectome when compared to the conventional disconnectome (see supplementary figure 2 for a side-by-side comparison of the two spaces). Thus, this suggests a greater differentiation for deep-disconnectomes derived from similar stroke lesions. Reversely, the distance between clusters is smaller for the deep-disconnectome than in the conventional disconnectome framework suggesting a more uniform spread within the morphospace. These differences may have improved the segregation between similar profiles of white-matter damage (i.e., in the same cluster) and, accordingly, would have led to better modeling of fine differences within the same neuropsychological assessment (Figure 2c).

## 3. Discussion

To our best knowledge, a deep learning model has not been used so far to obtain disconnectome maps. The deep learning 3D U-Net model we trained can generate accurate maps of brain dysconnectivity – deep-disconnectomes – derived from a simple binary lesion mask. These generated disconnectivity maps have a clinical predictive power toward long-term impairment following a stroke that is even higher than conventionally computed disconnectome maps. The validation of the deep-disconnectomes’ capacity to accurately predict neuropsychological scores demonstrate its potential to be released as an AI-driven tool for clinical applications.

Although our deep learning model generates disconnectome-like images, it does not capture exactly the same patterns as the conventionally computed disconnectomes. It can be speculated that the 3D U-Net model is potentially getting rid of noise while producing deep-disconnectomes, allowing for more concise information to be forwarded into the UMAP framework, consequently yielding more accurate predictions. Investigating this question, we conducted a paired t-test comparing conventional disconnectomes with their corresponding (derived from the same binary lesion mask) deep-disconnectomes. The results confirmed our speculation, revealing that deep-disconnectomes emphasize the core white-matter structures while exhibiting comparatively lower capacity in capturing short, more detailed, U-shaped fibers (Figure 1a). Furthermore, we compared the average Euclidean distances of deep-disconnectomes and conventional disconnectomes to all the other points in their respective morphospace (Figure 2b). We detected that deep-disconnectomes are better segregated within clusters of near data points. This means that deep-disconnectomes with small Euclidean distances (showing similar disconnectivity patterns) are dissociated better than conventional (BCBTookit) disconnectomes. On the contrary, deep-disconnectomes with large Euclidean distances (showing completely different disconnectivity patterns) are less different than conventional disconnectomes (which might not impact the predictive performances as these disconnectivity patterns are already well segregated). The difference between the deep-disconnectome and the conventional disconnectome cannot be driven by the ‘average template disconnectome’ used as a second input to the 3D U-Net model. Since this template is a fixed parameter of the model, the sole variability of deep-disconnectomes originates from individual lesion masks. Generating a deep-disconnectome without the ‘average template disconnectome’ did not produce sufficient results since the amount of information from the binary lesion mask was not enough to train a deep learning model, applying the parameters we used. This is a common challenge that related studies encountered in the past. Augmenting the input information in a 3D U-Net model with additional data sources or other contextual information has been shown to improve the outcomes in various medical imaging applications. Myronenko et al. (2019) demonstrated that the fusion of multiple modalities (T1, T2, FLAIR images) helped capture diverse tumor characteristics, improving the accuracy of the segmentation results of the model. Another example of a multi-model fusion approach was shown by Guasch and colleagues (2020), who developed a method inspired by geophysics that generates accurate images of the brain applying full-wave inversion. This, however, required their full-wave inversion model to contain an a priori acquired model of the skull, without which it would not have converged to a legitimate solution. While in this exemplary case, an individual skull model needs to be provided in order to achieve satisfactory results, in the present case of obtaining deep-disconnectomes, we only need one single ‘average template disconnectome’, which can be used as a constant for the model input. Providing this information alongside the lesion mask through two channels in the 3D U-Net model allows us to obtain individual predictions of the brain disconnectivity much faster than computing such maps using conventional methods, e.g., whole brain tractography.

While producing deep-disconnectomes with a trained AI model makes obtaining the otherwise processing-heavy data faster, the potential of its application goes beyond the advantage of gaining time. They outperform the conventional disconnectomes when applied in a recent framework for neuropsychological score predictions. Even though the obtained deep-disconnectomes overlapped with the conventional disconnectome maps reaching an average accuracy of R^2^=0.82 (Figure 1a), the captured information provides valuable insights into a patient’s disconnection pattern and could be used for further follow-up analyses and predictions. For instance, to evaluate the predicted deep-disconnectomes on a use case, we applied the framework of Talozzi et al. (2023) to predict neuropsychological scores for 86 tests covering different categories of cognitive and behavioral outcomes one year following a stroke. Incidentally, the prediction accuracy of neuropsychological scores using deep-disconnectomes exceeded the original predictions that used conventional disconnectome maps (Figure 2b). This result confirms that deep-disconnectomes can be used for such analyses. Although the ground truth information is not being reproduced with 100% accuracy, the deep learning 3D U-Net model outputs simplified disconnectomes from a binary lesion mask from which seemingly the UMAP algorithm for dimensionality reduction can extract features in a more precise manner. This added precision led to better regression coefficients and, consequently, to better predictions for clinical test outcomes.

Alongside the impressive performance of the simple 3D U-Net model comes the benefit of time and computational capacity needed to obtain a deep-disconnectome from a lesion mask, making it feasible and practical at the clinical level. This is a common advantage of deep-learning methods, which require a time-demanding training phase with numerous data, but have the tremendous gain of being fast when applied to new instances: a perfect fit for precision medicine suitable for clinical implementation. Using a trained deep-learning model requires the operator to have less than 10 MB of free space on their device, and a deep-disconnectome can be outputted within less than one second. This is an improvement in comparison to the current ways of computing brain disconnections using, e.g., toolkits such as the BCBToolkit (Foulon et al., 2018), the Network Modification (NeMo) Tool Lite (Kuceyeski et al., 2013), or the Lesion Quantification Toolkit (LQT) (Griffis et al., 2021). Usually, programs that are able to compute brain disconnectomes require at least 1,5 GB up to 10 GB of free memory space, taking approximately 720 times longer to create an output (0.642 seconds for the deep-disconnectome in comparison to 480 seconds for the conventional disconnectome). Moreover, the operator needs to follow precise pipelines when reconstructing conventional disconnectomes. The sequence of numerous command steps often leaves room for the erroneous setting of parameters (e.g., the threshold used) or software misuse (e.g., wrong templates for neuroimaging). Not only does the trained 3D U-Net offer a faster and more reliable way of obtaining information about brain disconnectivity, but it also yields more accurate predictions of individual clinical scores.

Finally, a deep learning model that creates disconnectomes based on minimal individual information circumvents the step of downloading software, makes obtaining disconnectivity data more accessible, and, by that, consequently speeds up scientific progress. However, while the integration of deep learning into various domains holds immense potential, its application still faces constraints. One prevailing limitation, which also pertains to our study, is the issue of generalizability. The efficacy of an algorithm heavily relies on the quality and representativeness of the training data it receives (Taylor & Nitschke, 2018). Consequently, ensuring a diverse and comprehensive dataset becomes crucial for maximizing the model’s ability to generalize effectively. Within the realm of medical diagnosis, this challenge is particularly pronounced, exacerbated by biases and stringent privacy policies that restrict data availability as well as typical bias in the distribution of clinical data (Gianfrancesco et al., 2018). However, in the specific context of disconnectomes, we are fortunate to have addressed this limitation by training our model on a broad sample of diverse synthetically generated lesions. Leveraging the capabilities of the BCBToolkit, we can generate an abundance of training data by computing disconnectomes. This unique advantage augments our framework and unlocks novel opportunities for further exploration. The deep-disconnectome is theoretically able to capture the pattern of disconnection for any type of focal lesion (such as focal multiple sclerosis lesions, brain tumors, and post-traumatic lesions). In future work, we can extend the model’s training to include alternative focal lesions and examine the correlation between deep-disconnectome outcomes and associated neuropsychological manifestations. Incorporating actual patient data with alternative focal lesions into our deep-learning model would enhance its overall generalizability and establish a more comprehensive foundation for predicting neuropsychological scores following brain damage. Despite the notable time-saving and precise outcomes achieved by AI models, there remains a prevailing mistrust among individuals regarding their reliability in healthcare diagnoses. While statistics demonstrate the better performance of AI models and the number of FDA-approved AI algorithms is steadily growing, concerns stem from the inherent opacity of these “black box” models, which oftentimes do not provide sufficient explanations for their results (Bzdok, Krzywinski & Altman, 2017). This lack of transparency undermines trust in their capabilities. With our framework for predicting stroke outcomes, we mitigate this issue by creating the deep-disconnectomes, which serve as a crucial intermediary, providing a direct connection between the structure and function of the brain and aiding in the interpretation of the predictions rather than predicting neuropsychological scores from lesions directly.

Overall, the achievement of accurately predicting deep-disconnectomes from binary lesion masks represents a significant initial milestone. This not only enriches the realm of neuroscience but also fosters the convergence of computer science with other disciplines, thereby pioneering advancements. In spite of this progress, the fields of neuroscience and computer science must collectively surmount the aforementioned challenges. Then, AI-driven models can more seamlessly find practical applications in clinical settings, enhance efficiency within healthcare systems, and ultimately contribute to an improved quality of life for individuals.

## 4. Method

### 4.1 Datasets for model training and testing

Our deep-learning framework was trained on a dataset of N=1333 synthetic lesions and their corresponding disconnectomes (Dataset 1) and subsequently tested for real stroke lesions (Dataset 2). The synthetic stroke lesions were initially computed by Thiebaut de Schotten, Foulon, and Nachev (2020) to assess the spatial distribution of stroke impact on cognition. For that, they produced more than one million synthetic stroke lesion masks to sample out 1333 lesions matching in hemispheric lateralization and voxel size with natural lesions but randomly distributed in the brain. The probability of disconnectivity for each voxel in the brain was calculated with the BCBToolkit (Foulon et al., 2018), and the obtained disconnectomes were used as ground truth to train our model. Using synthetic lesions had two main benefits: Firstly, we circumvented the eventual problem of insufficient training data since we could produce any desired number of synthetic lesion-disconnectome pairs. Secondly, the diversity of the training data increased, theoretically also increasing the generalizability of the trained model. During the training of our 3D U-Net, we used a split 80%/20% of synthetic data (Dataset 1) for, respectively, training and validation.

Subsequently, we used the N=1333 lesion masks segmented from patients recruited by the University College London Hospitals (Dataset 2) to test our trained deep-learning model. This patient cohort was recruited as part of a study approved by the West London and GTAC Research Ethics Committee. MRI scans (1.5 T and 3 T) of patients with acute ischemic stroke were acquired across several scanners between 1-2 weeks following the stroke (Xu et al., 2018). Patients (56% male) were, on average, 64 ± 16 years old (age range: 18–97 years). For more details on the used datasets, see Supplementary Material Table 1.

### 4.2 Deep learning 3D U-Net algorithm

We implemented a 3D U-Net network for predicting individual deep-disconnectomes from binary lesion maps (Figure 3a). This model can be considered as a series of encoding and decoding phases. The former learns underlying, high-level patterns from data, and the latter decodes the previous information into a desired format. Our encoder architecture consisted of three convolutional blocks, whereas the decoder transposed convolutions in three upconvolutional blocks to reconstruct the original image resolution. The encoder modules’ convolutional blocks consisted of two 3D convolutional layers with batch normalization and Leaky Rectified Linear Unit (ReLu) activation functions, followed by a max-pooling layer with a dropout rate of 0.5. In the encoding phase, the number of filters in each block is doubled (upsampling with factor 2) with respect to the previous block being gradually increased from 16 to 64 in order to increase the capacity of the model to learn more complex features. At the same time, the spatial resolution is reduced by half with each max-pool operation. In the decoder module, the upconvolutional blocks combine the output of the previous block with the corresponding output from the contracting path. Each of the upconvolutional blocks consists of an upsampling layer (using trilinear interpolation), a concatenation operation, a 3D convolutional layer with batch normalization, and a ReLu activation function. In convolutional layers, the size of all convolutional kernels is 3 × 3 × 3. For all max-pooling layers, the pooling size is 2 × 2 × 2 with stride= 2. The number of filters within each of the three upconvolutional blocks is 192, 96, and 48 filters, respectively. The spatial resolution of the output is doubled with each upsampling operation.

The model input consists of two 3D channels of the volume size 96×96×96 (voxels at 2mm^3). The first channel contains the binary stroke lesion, and the second channel is the average disconnectome image computed from 1333 synthetic disconnectomes using *fslmaths* as part of the FSL package (https://fsl.fmrib.ox.ac.uk/fsl/fslwiki; Jenkinson et al., 2012; Smith et al., 2004). The average disconnectome template was kept constant for every training and testing sample. During the data loading procedure, the images (91×109×91) undergo a size transformation containing interpolation to ensure equal dimensions of the images. In the evaluation, this transformation is reversed in order to compare the output of the model to the ground truth. The model obtains its final output by a 3D convolutional layer which is applied with a sigmoid activation function that maps the network output to the intensity range of the conventional disconnectome images (i.e., between 0 and 1, representing the probability of a voxel to be disconnected). Voxels outside of an a priori calculated brain mask based on the 2mm MNI152 T1 (Mazziotta et al., 2001) were excluded from model training, evaluation, and predictions. Local computational resources allowed for training with a batch size of four and a learning rate of 0.0003. During the training, the model updated its weights based on the sum of the MSE (Mean Squared Error) and MAE (Mean Absolute Error) loss functions. While the MSE loss measures the average squared difference between the predicted output and the target output, penalizing larger errors more heavily, the MAE loss measures the average absolute difference, being less sensitive to outliers. Combining these losses strikes a balance for penalizing errors, allowing to capture fine details, as well as the overall structure in the predicted outcome (Botchkarev, 2019). To minimize the loss, the model network weights were iteratively updated with a weight decay of 0.0001 using the Adam optimizer. The model training was programmed for a maximum of 300 epochs on an NVIDIA Quadro P4000 GPU (graphics memory of 8GB), with an implementation of stopping the training earlier (early stop) once the validation loss stopped decreasing for 50 epochs.

**Figure 3:**
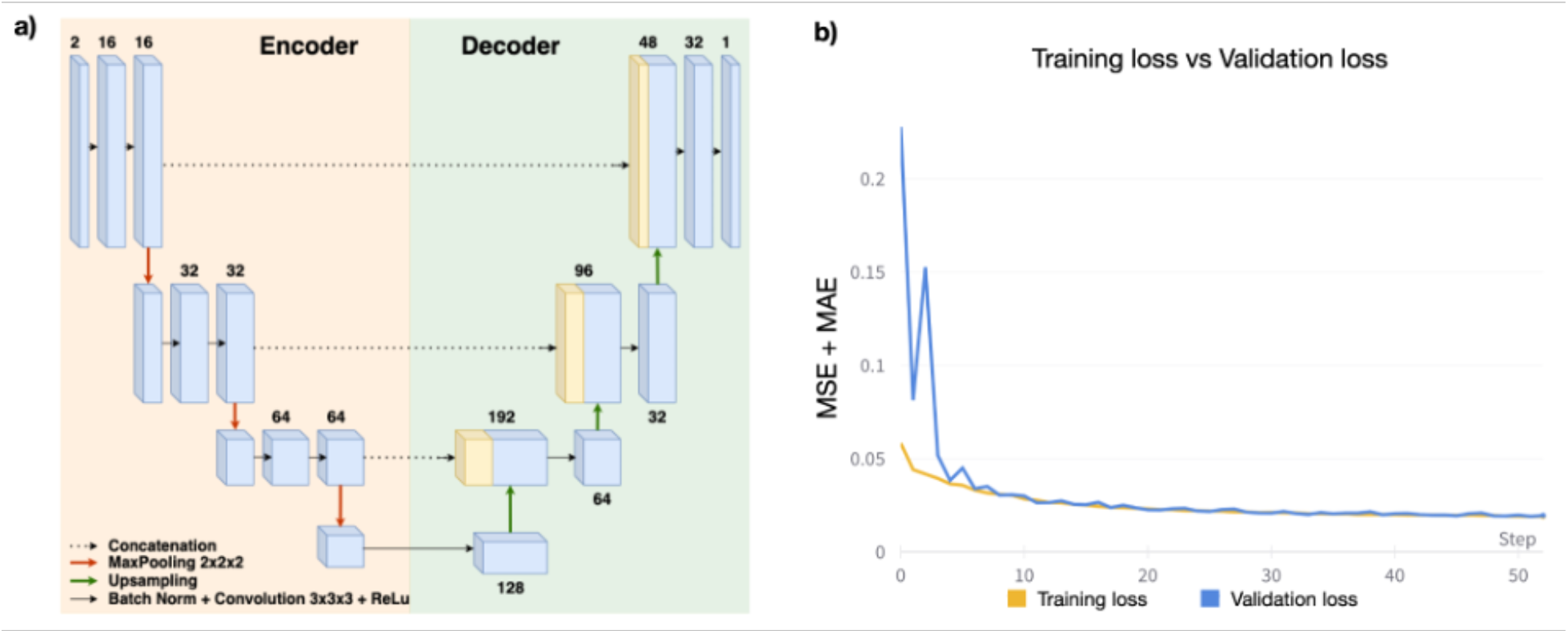
(a) Architecture of the 3D U-Net used for deep-disconnectome predictions. The number above/below each block is the number of the used channels. (b) Training and validation loss over the time course of the first 50 epochs was evaluated by adding the Mean Squared Error (MSE) and Mean Absolute Error (MAE).

### 4.3 Deep-disconnectomes and conventional disconnectomes comparison

To assess the level of similarities between deep-disconnectome and conventional disconnectomes, we first produced a Pearson R^2^ revealing the percentage of the variance of disconnection reproduced by the 3D U-Net (i.e., the goodness of fit).

To explore any systematic differences in terms of disconnected fibers, we ran a voxelwise paired t-test for dataset 2, comparing the conventional disconnectomes and their corresponding deep-disconnectomes. To circumvent any differences in the ranges of the intensity values (i.e., deep-disconnectomes typically do not produce 0 or 1 values), both disconnectomes were binarized using a threshold of 0.0001 (Figure 1b). Subsequently, the embedding properties were compared across the two disconnectome methods. For the comparison of the coordinates in the disconnectome morphospace, we computed the average Euclidean distances for 139 coordinate pairs (corresponding to the 119 data points of dataset 3, and 20 data points of dataset 4) across deep-disconnectomes and conventional disconnectomes. This comparison allowed us to explore the relative distances between data points in the embedding space. Subsequently, we visualized the results in a Bland Altman plot showing the differences in morphospace distances between conventional and deep-disconnectomes. In this visualization, a negative difference indicates a bigger distance between the data points, while a positive distance indicates a smaller distance between the data points. By comparing the distribution of the data points, we gain insights into the differences between the two types of disconnectomes that can be relevant to explain differences in the prediction of neuropsychological scores.

### 4.4 Prediction of neuropsychological scores one year after stroke

Achieving clinical endpoints is the main goal of our investigation of stroke disconnections. In order to test and compare the potential in clinical score prediction, we compared the deep-disconnectomes prediction and the conventional disconnectomes performances.

For this purpose, a third dataset was used (Dataset 3, N=119 stroke survivors). Neuropsychological score predictions were performed on datasets 3 and 4, recruited at the School of Medicine at Washington University in St. Louis (Corbetta et al., 2015). N=86 scores were collected, spanning from motor and language abilities to visuospatial, memory, and sickness reports. Details of the acquired scores can be found in Talozzi et al. (2023). The peculiarity of these datasets is to have neuroimaging within 2 weeks from the stroke event, whereas neuropsychological scores were evaluated on average one year later hence allowing to test the long-term predictive power of acute neuroimaging. This information could be key for informing behavioral therapies. To test the predictive quality of the deep-disconnectomes obtained from the 3D U-Net model, we replicated the analysis and validation procedures of Talozzi et al.’s (2023) work. The authors were able to accurately predict neuropsychological scores 1 year after stroke by creating a latent disconnectome morphospace using the UMAP algorithm for dimensionality reduction (McInnes, Healy & Melville, 2018). First, 1333 disconnectomes of dataset 2 (corresponding to the ‘real’ lesions used in the presented testing procedure of our 3D U-Net) were used to compute the 2D UMAP morphospace based on the standard computation of disconnectomes through the BCBToolkit software. Following this step, disconnectome maps from dataset 3 (n = 119) were imported into the UMAP morphospace, and their localization was statistically correlated to the individual neuropsychological scores on the N=86 behavioral and cognitive scores. To handle multiple clusters of voxels that survived the threshold for medium effect size (R>0.2), a principal component analysis (PCA) was run to capture the greatest source of patient coordinate variability. Based on the first three PCA components, a multiple regression model (Python *scikit-learn* package) was evaluated to predict each neuropsychological score. For more detailed information on the methods, see Talozzi et al. (2023). Note that the datasets used in the present paper are the same as in Talozzi et al. (2023), the only difference being the method of how the disconnectomes were obtained: in Talozzi’s work using BCBToolkit software, here using a trained 3D U-Net model. Hence, conventional disconnectomes and deep-disconnectomes are systematically comparable.

## Code Availability

The trained model, the codes used, and exemplary files can be found at https://github.com/annamatsulevits/deep-disconnectome.

## Acknowledgment

This work was supported by the European Union’s Horizon 2020 research and innovation programme under European Research Council (ERC) Consolidator grant agreement no. 818521 (M.T.d.S., DISCONNECTOME); University of Bordeaux’s IdEx “Investments for the Future” program RRI “IMPACT,” which received financial support from the French government.

