## Supplementary Information for "Deep Learning disconnectomes to accelerate and improve long-term predictions for post-stroke symptoms"

**Supplementary Material**

**Table 1:** Information on the used datasets

| | Dataset 1 | Dataset 2 | Dataset 3 | Dataset 4 |
|---|---|---|---|---|
| Recruitment site | Synthetically produced (Thiebaut de Schotten, Foulon, and Nachev (2020)) | University College London Hospitals, London (UK) | School of Medicine of the Washington University, St. Louis (USA) | School of Medicine of the Washington University, St. Louis (USA) |
| Conducted analyses | 3D U-Net training | 3D U-Net testing, deep-disconnectome production, and UMAP space creation | UMAP training, 86 neuropsychological scores | out-of-sample testing of predictions, 86 neuropsychological scores |
| **Demographics** | | | | |
| N | 1333 synthetic lesions | 1333 real lesions | 119 | 20 |
| Males/females, *n* | NA | 748/585 | 65/54 | 12/8 |
| Age, years | NA | 64 ± 16 (18–97) | 54 ± 11 (19–83) | 58 ± 12 (34–95) |
| Education, years | NA | NA | 13.2 ± 2.5 (5–20) | 13.7 ± 2.6 (9–19) |
| Right-handed/left-handed, *n* | NA | NA | 109/10 | 17/3 |
| **MRI** | | | | |
| Chronology | NA | 1–2 weeks after stroke | 14 ± 8 days after stroke | 13 ± 4 days after stroke |
| Dominant lesion site | 44% right hemisphere 56% left hemisphere | 44% right hemisphere 56% left hemisphere | 46% right hemisphere 54% left hemisphere | 40% right hemisphere 60% left hemisphere |
| **Neuropsychological assessment** | | | | |
| Delay after stroke onset, days | NA | NA | 393 ± 56 | 385 ± 22 |
| Delay after MRI scan, days | NA | NA | 379 ± 57 | 373 ± 22 |

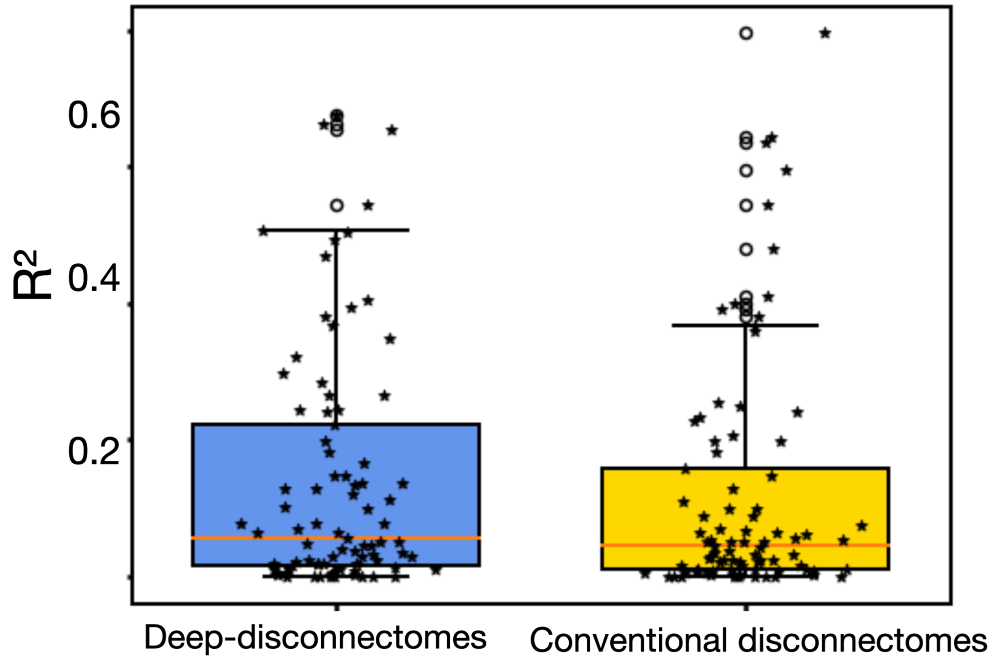


**Figure 1:** The boxplot shows all the R² for the predictions of the validation set (N=20) across N=86 neuropsychological scores for the framework using deep-disconnectomes compared to the framework using disconnectomes obtained by BCBToolkit. The boxes represent the quartiles, the whiskers indicate the distribution, and the outliers are marked as white dots. Inside the boxes, the median is visualized by the red line. The *P*-value obtained from a paired *t*-test (2-tails): *P*=0.241 shows no significant difference in the prediction power for this out-of-sample cohort.

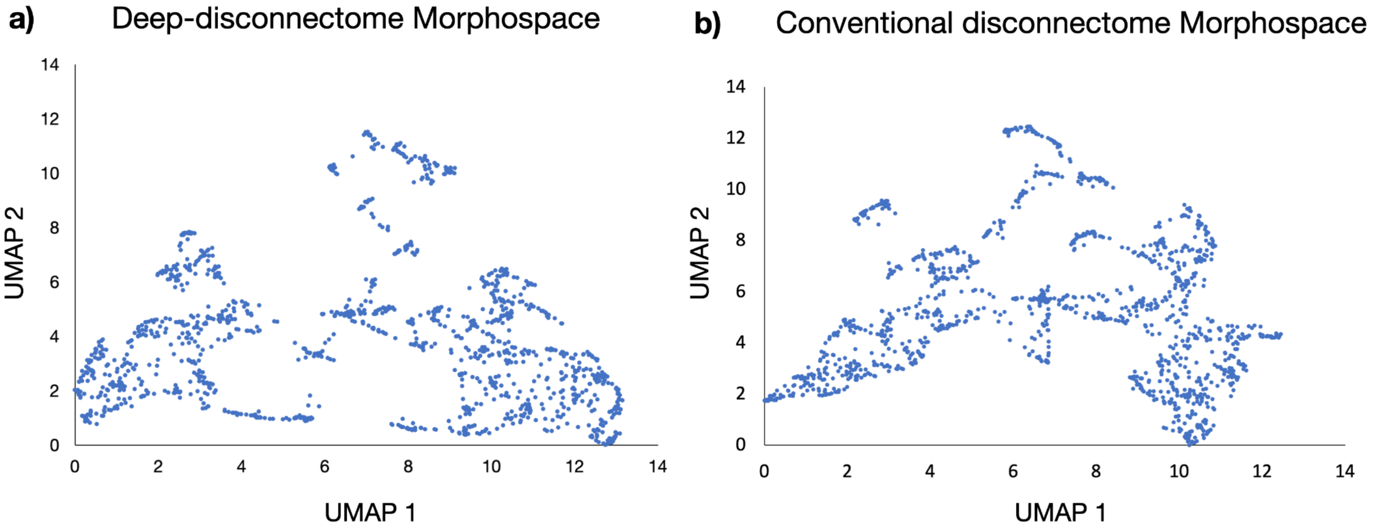


**Figure 2:** Comparison of the morphospaces obtained using different disconnectomes. (a) Deep-disconnectome morphospace. (b) Conventional disconnectome morphospace
